# Transposable elements as tissue-specific enhancers in cancers of endodermal lineage

**DOI:** 10.1101/2022.12.16.520732

**Authors:** Konsta Karttunen, Divyesh Patel, Jihan Xia, Liangru Fei, Kimmo Palin, Lauri Aaltonen, Biswajyoti Sahu

## Abstract

Transposable elements (TE) are repetitive genomic elements that harbor binding sites for human transcription factors (TF). A regulatory role for TEs has been suggested in embryonal development and diseases such as cancer but systematic investigation of their functions has been limited by their widespread silencing in the genome. Here, we have utilized unbiased massively parallel reporter assay data using whole human genome library to identify TEs with functional enhancer activity in two human cancer types of endodermal lineage, colorectal and liver cancers. We show that the identified TE enhancers are characterized by genomic features associated with active enhancers, such as epigenetic marks and TF binding. Importantly, we identified distinct TE subfamilies that function as tissue-specific enhancers, namely MER11- and LTR12-elements in colon and liver cancers, respectively. These elements are bound by distinct TFs in each cell type, and they have predicted associations to differentially expressed genes. In conclusion, these data demonstrate how different cancer types can utilize distinct TEs as tissue-specific enhancers, paving the way for comprehensive understanding of the role of TEs as bona fide enhancers in the cancer genomes.

## Introduction

Around half of the human genome consists of sequences that originate from TE insertions. The advent of whole genome sequencing technologies has revealed the major contribution of TEs to the evolution, size and regulatory functions of eukaryotic genomes[1]. TEs harbor *cis*-regulatory sequences for human TFs and can thus contribute to the human gene regulatory elements such as promoters and enhancers[2, 3]. TEs have been shown to have regulatory functions during development[4], and their role in complex genetic diseases such as cancer is also becoming more evident[5]. However, to what extent TEs contribute to gene regulatory functions in different cancer types is still poorly understood.

Most of the TEs in the human genome have been immobilized due to truncations and mutations that accumulate during evolution and majority of TEs remain epigenetically silenced in normal somatic cells. However, there are ∼100 copies of long interspersed nuclear element 1 (LINE-1; L1) that are capable of retrotransposition, *i*.*e*., inserting themselves to new genomic loci[6]. Accumulating evidence from whole genome sequencing studies indicates that the L1 elements can be widely activated in cancer[7-10] and different cancer types show distinct rates for somatic retrotransposition[11, 12]. Aberrant TE insertions can contribute to tumorigenesis by inducing genomic rearrangements that can lead to deletion of tumor-suppressor genes or amplification of oncogenes[11]. Importantly, the retrotransposition-incapable TEs can also play a role in tumorigenic processes by providing a large repository of potential regulatory elements that can be repurposed for transcriptional control of human genes[13]. However, since the activation of the retrotransposition-incapable TEs can occur for example through destabilization of their epigenetic silencing, their activation cannot be detected in whole genome sequencing data and thus their functional significance in cancer cells has remained largely elusive. Few studies have reported activation of proto-oncogenes by derepressed long terminal repeat (LTR)-elements at their promoters[14-16], but systematic functional studies and epigenetic profiling are necessary for comprehensive understanding of the regulatory activity of TEs in the cancer genomes.

Here, we have leveraged on the unbiased and genome-wide episomal enhancer activity measurements from an ultracomplex whole human genome library together with genome-wide epigenetic profiling to characterize TE-derived enhancers in two distinct cancer types, colorectal and hepatocellular carcinomas. We show that the functional activity is similar for a subset of TEs but, importantly, there are distinct TE subfamilies that are activated by different TFs in a highly tissue-specific manner, suggesting that these TEs can act as tissue-specific enhancers resulting in differential gene regulatory activity in human cancers.

## Results

### Long terminal repeats are enriched within active enhancers identified by STARR-seq

To measure the functional enhancer activity of TEs in two endodermal origin cancers, namely colon and liver cancers, we utilized our publicly available massively parallel reporter assay data from the genomic STARR-seq experiments in GP5d and HepG2 cell lines, respectively[17]. We then characterized the enhancer properties of the TEs by combining STARR-seq with epigenetic and regulatory data from the same cell lines, such as ATAC-seq for chromatin accessibility, ChIP-seq for epigenetic marks and TF binding, and nuclear run-on data for detecting the nascent transcripts (**Fig. 1a**). In the STARR-seq datasets utilized here, the whole human genomic DNA was cloned into the 3’-UTR of the reporter gene driven by a weak minimal promoter to measure enhancer activity with a ∼1.5 bp resolution[17]. We mapped the data to human genome (hg38) and identified the active enhancers by peak calling against plasmid input, resulting in 15,390 peaks for GP5d and 11,951 peaks for HepG2 (**Supplementary table 1 and 2**). Over half of the active enhancer peaks in both GP5d and HepG2 overlap with at least one TE (**Fig. 1b**). Analysis of different TE classes [LINE, short interspersed nuclear elements (SINE), LTR, and DNA elements] revealed that the LTR elements were significantly overrepresented within the active enhancer peaks in both cell lines (p < 2.2×10^−16^; **Fig. 1c**). Many LTR subfamilies were highly occupied with STARR-seq peaks with up to 40% of their genomic copies overlapping a peak summit (**Extended Data Fig. 1a**). Previously, it has been shown that evolutionarily young LTRs are enriched within the open chromatin more frequently compared to older LTRs, suggesting that younger LTRs are more active in the genome[18]. This was shown to be independent of their sequence features, such as mappability of TE subfamilies[18]. In agreement with this, we found primate-specific TEs being overrepresented within the active enhancers in both cell lines (**Fig. 1d**). Taken together, our results show that we can detect enhancer activity from the LTR elements in both colon and liver cancer cells in the STARR-seq assay.

**Figure 1.**
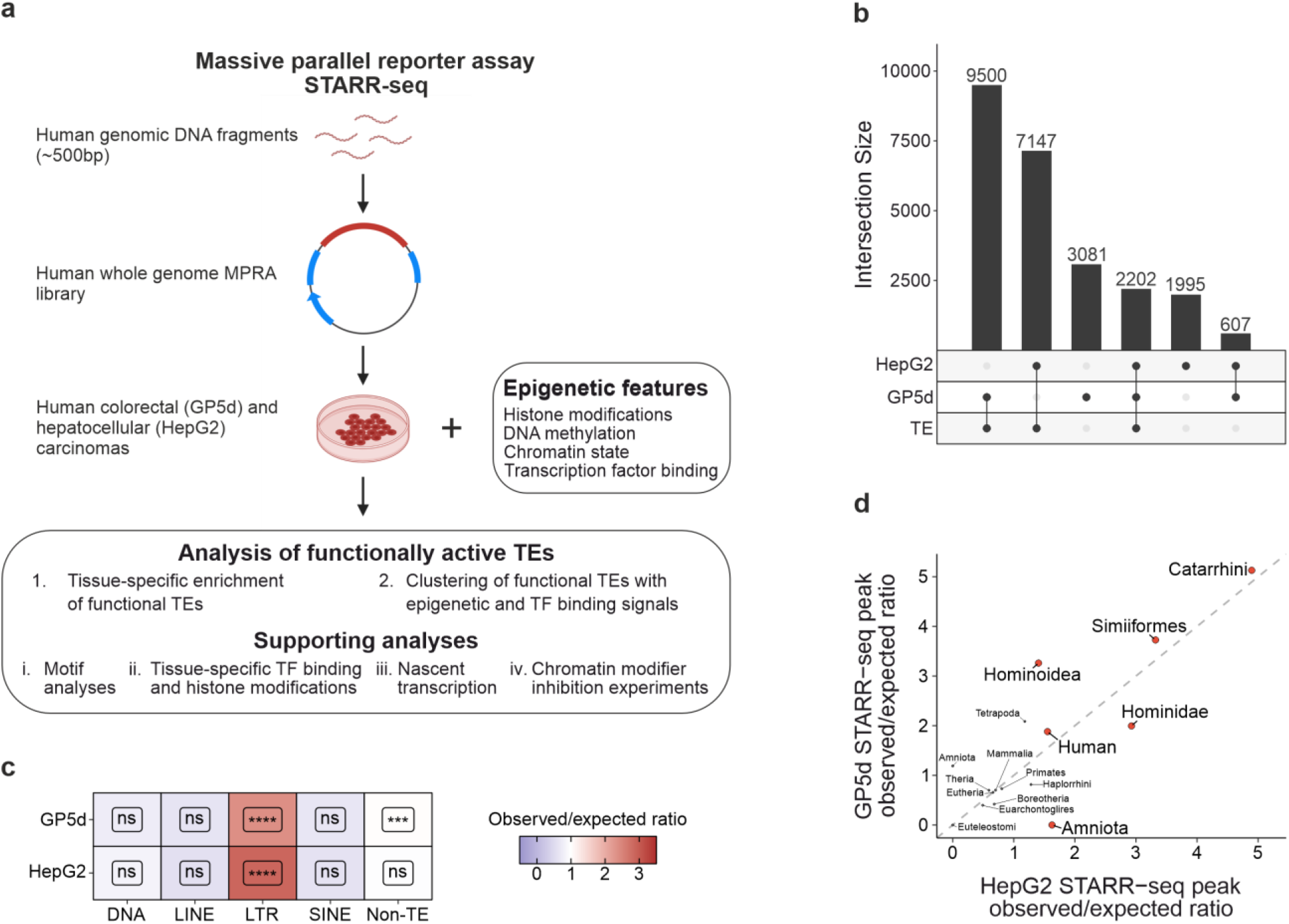
Long terminal repeats are enriched within active enhancers identified by STARR-seq. a) Schematic representation of the analysis pipeline. b) Upset plot for overlap analysis of STARR-seq peaks with TEs in GP5d and HepG2 cells with the number of peaks in each category indicated; total number of peaks 15,390 and 11,951 in GP5d and HepG2 cells, respectively. c) Ratio of observed vs. expected overlaps for GP5d and HepG2 STARR-seq peak summits with the major classes of TEs (DNA, LINE, LTR and SINE) and the non-TE genome. BH-adjusted one-sided binomial test FDR is shown for each class (Significance symbols: **** indicates p < 0.0001, *** p < 0.001, ** p < 0.01, * p < 0.05, ns = non-significant, p > 0.05). d) Enrichment of STARR-seq peak summits at TEs classified by lineage significant in both GP5d and HepG2 cells. TE subfamilies were grouped by their lineage of origin and the observed/expected ratio of STARR-seq peak summits at the TE lineage groups was calculated. TE lineages significant in both GP5d and HepG2 (BH-adjusted one-sided binomial test FDR < 0.01) are labeled in red, gray points are statistically insignificant.

### TEs are enriched for distinct epigenetic signatures and TF binding motifs

To characterize the properties of active enhancers in GP5d cells, unsupervised k-means clustering was performed for all genomic STARR-seq peaks along with the signal for open chromatin from ATAC-seq, ChIP-seq for histone modifications (H3K4me1, H3K9me3, H3K27ac, H327me3, and H3K36me3) and p53 binding (with and without 5-fluorouracil treatment) as well as CpG methylation called from long-read nanopore sequencing. Five clusters resulted in optimal clustering (**Extended Data Fig. 2a**), and each cluster showed distinct enrichment for genomic regulatory features (**Fig. 2a**). Clusters 1 and 2 with 2,367 and 897 peaks harbor the classical features of enhancers and promoters, respectively. Both show strong chromatin accessibility and a decrease in CpG methylation, H3K9me3 and H3K27me3 signals (**Fig. 2a**). Peaks in cluster 1 are enriched for H3K4me1 and H3K27ac histone modifications with a bimodal signal around the peak center that is characteristic to active enhancers, whereas peaks in cluster 2 show very high enrichment for promoter-specific H3K4me3 and less prominent central distribution of H3K27ac signal. In addition, the flanks of the cluster 2 peaks are enriched for H3K36me3 that marks the gene bodies, and peak annotation confirmed their association to promoters in the human genome (**Extended Data Fig. 2b**). Enrichment analysis for TF binding motifs revealed basic leucine zipper (bZIP) family motifs such as JUN/FOS within the peaks in Cluster 1 (**Fig. 2b**), supporting their canonical enhancer function, whereas Cluster 2 was enriched for KLF/SP motifs from bZIP family as well as zinc finger factors from the YY family (**Fig. 2b**), which are known to be promoter-specific factors[19]. Cluster 2 peaks had very little overlap with TEs (**Extended Data Fig. 2c**), consistent with previous findings of TE depletion in promoter regions[20]. Interestingly, however, two thirds of the peaks in Cluster 1 overlap with TEs (**Extended Data Fig. 2c**), speaking for their potential as functional enhancers from the TEs in the endogenous genomic context.

**Figure 2.**
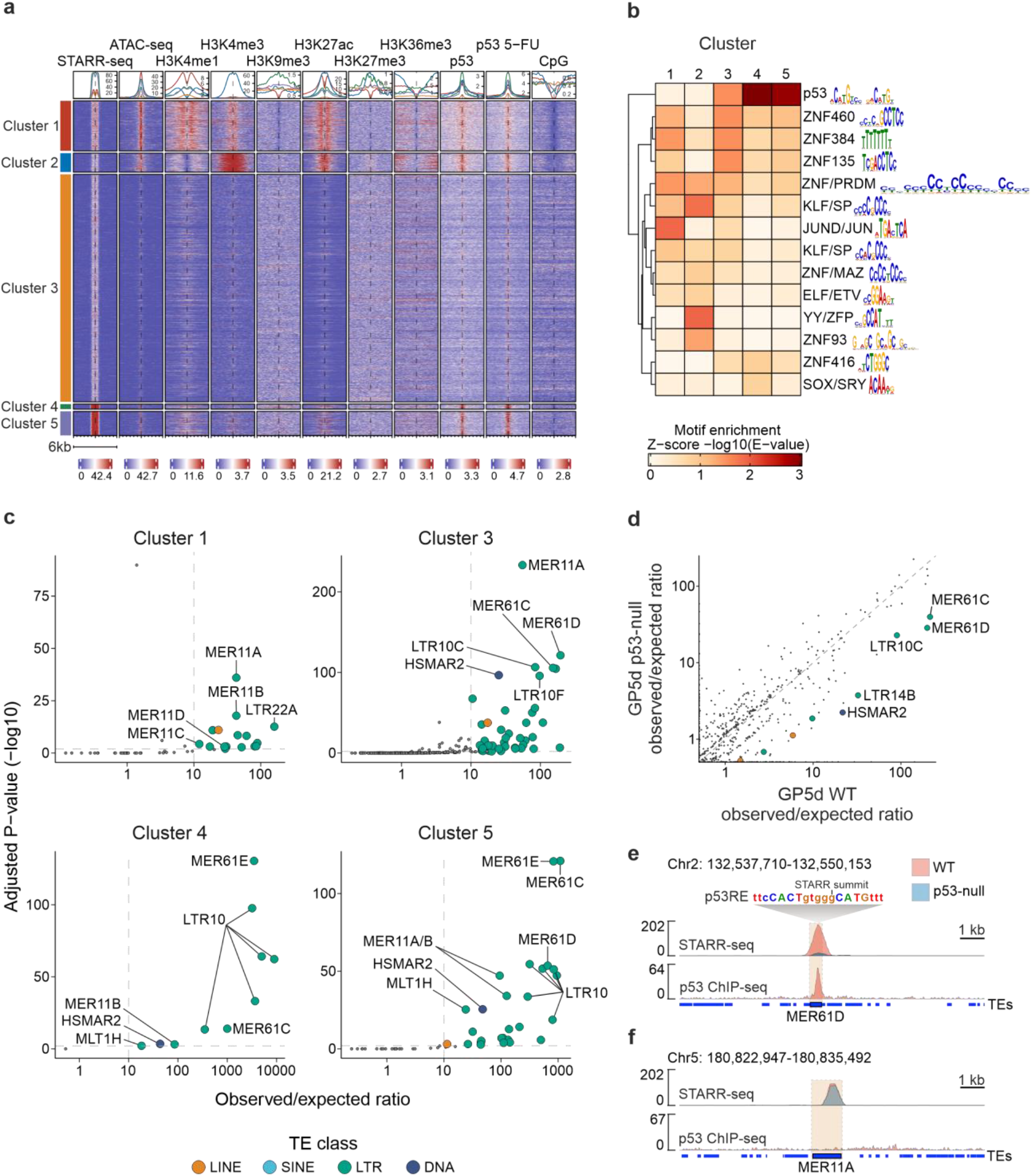
TEs are enriched for distinct epigenetic signatures and TF binding motifs in colon cancer cells. a) K-means clustering of STARR-seq peaks with open chromatin (ATAC-seq), ChIP-seq for epigenetic marks and transcription factor binding (H3K4me1, H3K4me3, H3K9me3, H3K27ac, H327me3, H3K36me3, p53 and 5-fluorouracil-treated p53), and CpG methylation (NaNOMe-seq). The signal is plotted ±3kb from peak summit. Five clusters was optimal for clustering as determined by elbow plotting (see **Extended Data Fig. 2a**). b) TF motif enrichment within each cluster from panel **a**. After performing the motif enrichment analysis for individual motifs, similar motifs were combined into motif clusters according to Vierstra, Lazar [26]. The representative TF family and motif are shown for each clustered motif. c) Enrichment of TE subfamilies within STARR-seq peaks of each cluster. Cluster 2 is omitted due to no significant enrichment. Observed/expected ratio on X-axis is calculated as the count of STARR-seq summits overlapping TE subfamilies divided by the mean count of 1000 repetitions of randomly shuffled STARR-seq peak summits overlapping TE subfamilies. Adjusted P-value on Y-axis is the –log10-transformed BH-adjusted binomial test FDR. Dashed line on Y-axis represents the significance limit (p = 0.01) and on X-axis the observed/expected ratio of 10 used as the limit of significance for STARR-seq peak summit overlap with TE subfamilies. d) Comparison of TE subfamily enrichment in STARR-seq peak summits between wild type (WT) and p53-null GP5d cells. Colored points mark differentially enriched TE subfamilies (One-sided Fisher’s exact test p < 0.01). Significant subfamilies with an observed/expected ratio of larger than 10 are labeled. e) Genome browser view of an example STARR-seq peak overlapping a MER61D insertion. The tracks for STARR-seq and p53 signals from the two cell lines are shown using the same scale and overlayed on top of each other, showing a loss of STARR-seq signal and the p53 binding at this MER61D locus due to p53 deletion. f) Genome browser view of an example STARR-seq peak overlapping a MER11A element, showing no p53 binding and equal STARR-seq signal in both WT and p53-null GP5d on the locus.

Cluster 3 with 10,741 peaks showed a major overlap with TEs but low STARR-seq signal and little enrichment for open chromatin or active histone marks (**Fig. 2a, Extended Data Fig. 2c**), suggesting that this cluster mostly comprises peaks that are repressed in the endogenous chromatin context but show enhancer activity in the episomal STARR-seq assay and resembling the cryptic enhancers described previously[17]. Motif analysis revealed moderate enrichment for p53 domain as well as zinc finger factor (ZNF) motifs that are mostly absent from other clusters (**Fig. 2b**), suggesting that repression of the TEs in this cluster is mediated *in vivo* via known TE suppressors, p53[21] and Krüppel-associated box domain zinc fingers (KRAB-ZNFs)[22]. However, the strongest enrichment for p53 domain motif was observed within Clusters 4 (n = 242) and 5 (n = 1143) (**Fig. 2b**). These clusters also showed the strongest STARR-seq signal indicating strong enhancer activity (**Fig. 2a**), and the majority of the peaks were associated with a TE (**Extended Data Fig. 2c**). The enrichment of active histone marks at the genomic loci corresponding to Clusters 4 and 5 was relatively low, but both showed strong p53 binding (**Fig. 2a**). Interestingly, Cluster 4 also showed a relatively high enrichment for H3K9me3 signal (**Fig. 2a**), an epigenetic mark that is known to be downregulated by p53 via the repression of lysine methyltransferase SUV39H1 activity[23]. This suggests that some of the peaks in Cluster 4 may be p53 targets that are active in the episomal STARR-seq assay but repressed *in vivo*.

Since the STARR-seq peaks within Clusters 1, 3, 4, and 5 showed major overlap with TEs (**Extended Data Fig. 2c**), we next analyzed enrichment of TEs at the subfamily level (**Fig. 2c**). In Cluster 4, almost all peaks overlap with a TE, which is reflected in the high relative enrichment of the subfamilies (**Fig. 2c, Extended Data Fig. 2c**). Peaks in Clusters 4 and 5 overlap with genomic p53 binding and we found that they are specifically enriched for TE subfamilies such as MER61E and LTR10 (**Fig. 2c**) that have been previously reported to harbor near-perfect p53 binding motifs[24]. The strong STARR-seq signal observed from these clusters is consistent with the previous reports that have demonstrated strong activity in a STARR-seq assay from p53-binding enhancers[17, 25].

The strongest enrichment for active enhancer marks was observed from peaks within Cluster Interestingly, Cluster 1 showed specific enrichment of all MER11 subfamilies, MER11A, MER11B, MER11C and MER11D (**Fig. 2c**), indicating that they may have an active role as enhancers in GP5d colon cancer cells. Cluster 3, on the other hand, harbors peaks that are mostly silenced in the endogenous genomic context. The TE subfamilies that were found to be enriched within this cluster were similar to all other clusters, including MER11, MER61 and LTR10 subfamilies (**Fig. 2c**), suggesting that the members of the same subfamilies can show different levels of enhancer activity and be under different epigenetic contexts in the same cell. Taken together, by integrating the unbiased enhancer activity data from episomal STARR-seq assay to epigenetic data at corresponding genomic loci, we have identified specific TE subfamilies that are either repressed or active enhancers in human colon cancer cells.

### p53 knockout reduces enrichment of specific TE subfamilies in GP5d cells

Due to the known p53-specificity of some TE subfamilies[24] that was corroborated by our findings of high enrichment of MER61 and LTR10 elements within Clusters 4 and 5, we set out to specifically study the role of p53 in TE-derived enhancer activity. For this, we analyzed previously published STARR-seq data from p53-null (p53-KO) GP5d cells, resulting in 13,349 active enhancer peaks in the absence of p53[17].We observed that depletion of p53 in GP5d cells led to a significant decrease in enrichment for nine TE subfamilies (p < 0.01; **Fig. 2d,e**), including MER61 and LTR10 subfamilies. Overall, the enrichment of TE subfamilies within the STARR-seq peaks correlated with the proportion of their genomic copies harboring p53 binding motifs (**Extended Data Fig. 2d**), and as expected, the TE subfamilies affected by p53 depletion were highly p53-specific: for example, 53.4% of MER61C copies harbor a p53-response element (p53RE)[24]. However, not all p53-specific TE subfamilies were affected by p53 depletion. For example, MER61E with 35.4% of its genomic copies harboring a p53 binding motif showed a reduction from 80-fold in WT to 57-fold in p53-null, but the effect was not statistically significant. This could be due to the redundancy of the p53 family members, suggesting that other members can bind and control these elements upon depletion of p53. In conclusion, our results indicate that p53 controls TEs in a highly specific manner, and it is not associated to all TE subfamilies, as shown for example for MER11A (**Fig. 2f**).

### STARR-seq reveals common and cell-type specific TE enhancers

To further interrogate if TE-derived enhancers are largely similar between cell types owing to their repetitive nature, we compared the enriched TE subfamilies for enhancer activity between GP5d and HepG2 cells. In GP5d cells, we identified 62 significantly enriched subfamilies within the STARR-seq peaks overlapping TEs (p-value < 0.01) (**Fig. 3a, Supplementary table 3**). Of these, 56 were from the LTR class, four from the LINE class and one each from the DNA and SINE class of TEs. This is concordant with a previous report showing that LTR elements in the human genome are enriched for TF binding sequences and often adapted to regulatory roles in the genome[27]. In HepG2 cells, the analysis of STARR-seq peaks overlapping TEs revealed a similar pattern, with 58 TE subfamilies significantly enriched (**Fig. 3a, Supplementary table 3**), majority of which (48) belong to the LTR class, five to the DNA class, four to the LINE class, and one to the SINE class. Of the significantly enriched TE subfamilies, 26 were common to both GP5d and HepG2 cells (p-value for enrichment < 0.01 in both, p-value for differential enrichment between cell lines > 0.01) (**Fig. 3a**, *left panel*; **Supplementary table 3**). The common subfamilies that showed the highest enrichment in both cell lines, such as MER61C, MER61E and LTR10B1, were highly specific for p53[24]. This was confirmed by motif analysis that showed high enrichment of p53 motifs within the common TE subfamilies (**Fig. 3b**). The observed/expected ratio of LTRs correlated in both cell lines with the percentage of genomic copies in a subfamily containing p53REs (**Extended Data Fig. 3a**; GP5d Pearson R = 0.8, p < 2.2e-16, HepG2 Pearson R = 0.71, p = 2.2e-16). In conclusion, these results indicate that p53-bound TE enhancers are conserved in cancer cell lines with wild-type p53.

**Figure 3.**
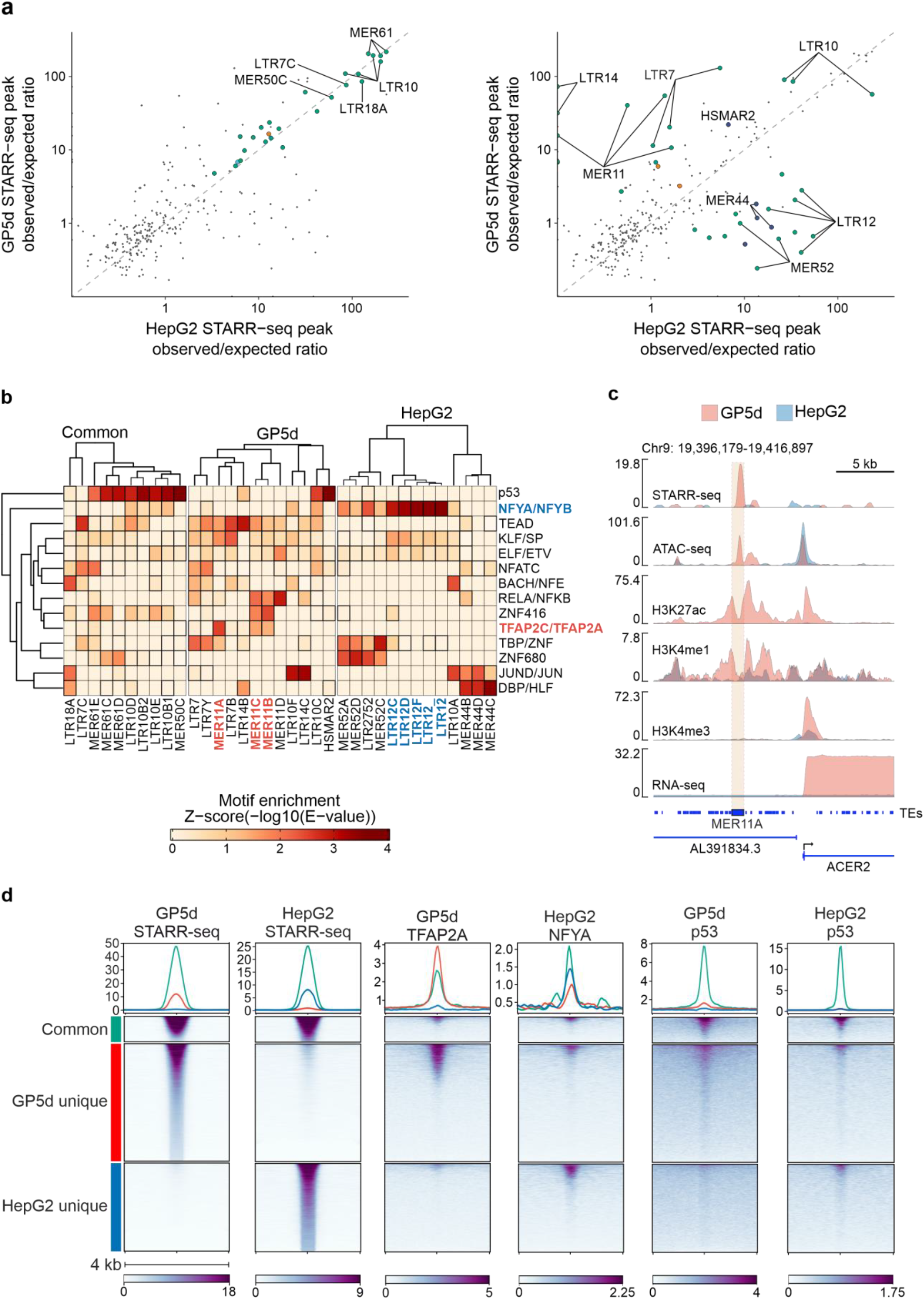
STARR-seq reveals common and cell-type specific differentially enriched TE subfamilies. a) Enrichment of TE subfamilies within STARR-seq peaks in GP5d and HepG2 cells. Only TE subfamilies with a minimum of five overlapping peak summits in GP5d or HepG2 are shown. In the left panel, the labels indicate the subfamilies that are significantly enriched in both cell lines (BH-adjusted two-sided binomial test FDR < 0.01, STARR-seq peak summit overlaps per subfamily >= 5) and not significantly differentially enriched between the cell lines (BH-adjusted two-sided Fisher’s exact test FDR > 0.01). Right panel indicates the differentially enriched TE subfamilies between the cell lines (BH-adjusted two-sided Fisher’s exact test FDR < 0.01, STARR-seq peak summit overlaps per subfamily >= 5). TE subfamilies are labeled as one group, e.g. MER11 group contains MER11A, MER11B, MER11C and MER11D subfamilies. **Supplementary table 3** lists all the enriched subfamilies. b) Motif enrichment for individual TE subfamilies. All significant common and differentially enriched TE subfamilies in GP5d and HepG2 (from **Fig. 3a**) were analyzed by taking the full sequences of the individual TEs with an overlapping STARR-seq peak and performing motif analysis for the sequences in each individual subfamily separately. The analysis was performed separately for common subfamilies as well as for subfamilies differentially enriched between GP5d and HepG2 cells. After motif analysis, the enriched motifs were grouped into clusters of similar binding motifs[26] (see Methods for details). The minimum E-value found for an individual TF for each motif cluster was plotted in the final figure. All of the found individual motif hits are listed in **Supplementary table 4**. c) Genome browser snapshot of a STARR-seq peak overlapping a MER11A element, showing STARR-seq and open chromatin signals as well as ChIP-seq track for canonical histone marks of active enhancers (H3K27ac, H3K4me1) in GP5d and HepG2 cells. Of note, the active enhancer marks as well as open chromatin is observed specifically in GP5d cells whereas the HepG2 cells show negligible signals at this locus. d) Heatmap showing signals for GP5d STARR-seq, HepG2 STARR-seq, TFAP2A and p53 ChIP-seq tracks from GP5d cells, and NFYA and p53 ChIP-seq tracks from HepG2 cells for three groups of STARR-seq peaks overlapping with TEs: shared between GP5d and HepG2 (Common, n = 2,202), unique to GP5d (n = 9,500), and unique to HepG2 (n = 7,147). Signal was plotted in ±2 kb flanks from the center of peaks.

Next, we analyzed cell type-specificity of TE enhancers and observed 18 TE subfamilies that were differentially enriched between GP5d and HepG2 cells (p-value for enrichment < 0.01 in at least one cell line, p-value for differential enrichment between cell lines < 0.01) (**Fig. 3a**, *right panel*; **Supplementary table 3**). On average, differentially enriched subfamilies were evolutionarily younger than the commonly enriched elements between the two cell lines (**Extended Data Fig. 3b**). Specifically, THE1, MER44, MER52 and LTR12 subfamilies were overrepresented in HepG2 cells and LTR14, MER11 and LTR7 subfamilies in GP5d cells (**Fig 3a,c**; **Supplementary table 3**). Interestingly, the differentially enriched TE subfamilies also showed over-representation of distinct TF motifs (**Fig. 3b**). For example, TFAP2 motifs were enriched within MER11 subfamilies in GP5d and NFY motifs within LTR12 subfamilies in HepG2 cells, suggesting that distinct TFs bind to TEs in a cell type-specific manner. To confirm that the TE enhancers identified from STARR-seq are bound by the TFs suggested by the motif analysis, we mapped the ChIP-seq data for TFAP2A in GP5d cells, for NFYA in HepG2 cells, and for p53 in both cell lines to the TE-overlapping STARR-seq peaks. In good agreement with the motif enrichment analysis, we observed p53 binding mostly at the common TE elements in both GP5d and HepG2 cells (**Fig. 3d**). TFAP2A was almost exclusively bound to GP5d-unique TEs in GP5d cells and NFYA preferentially bound to HepG2-unique TEs in HepG2 cells (**Fig. 3d**). Taken together, these results indicate that TEs show cell type-specific enhancer activity that is mediated by the binding of distinct TFs.

### Distinct TFs regulate cell type-specific transcriptionally active TE enhancers

To delineate TF binding dynamics at the TE subfamily level, we analyzed ChIP-seq signal at the specific LTR subfamilies that were overrepresented in distinct STARR-seq peak clusters in GP5d cells (*c*.*f*. **Fig. 2a, c**). As seen from the clustering, subfamilies within the MER family showed different epigenetic signals as MER11 elements associated with classical enhancer marks and MER61 elements with p53 binding. Analysis of ChIP-seq data at these loci revealed distinct TF occupancy: TFAP2A is highly enriched at MER11B (**Fig. 4a**) and MER11A (**Extended Data Fig. 4a**) loci, whereas MER61E and MER61C are highly specific for p53 binding (**Fig. 4b**; **Extended Data Fig. 4b**). Moreover, TF motif enrichment analysis suggested that HepG2-unique LTR12C elements are controlled by NFYA (*c*.*f*. **Fig. 2b**), and the analysis of NFYA ChIP-seq confirmed binding of NFYA at these loci in HepG2 cells (**Fig. 4c**). These results indicate that distinct TFs can bind specific TE subfamilies and utilize them as enhancers in a tissue-specific manner.

**Figure 4.**
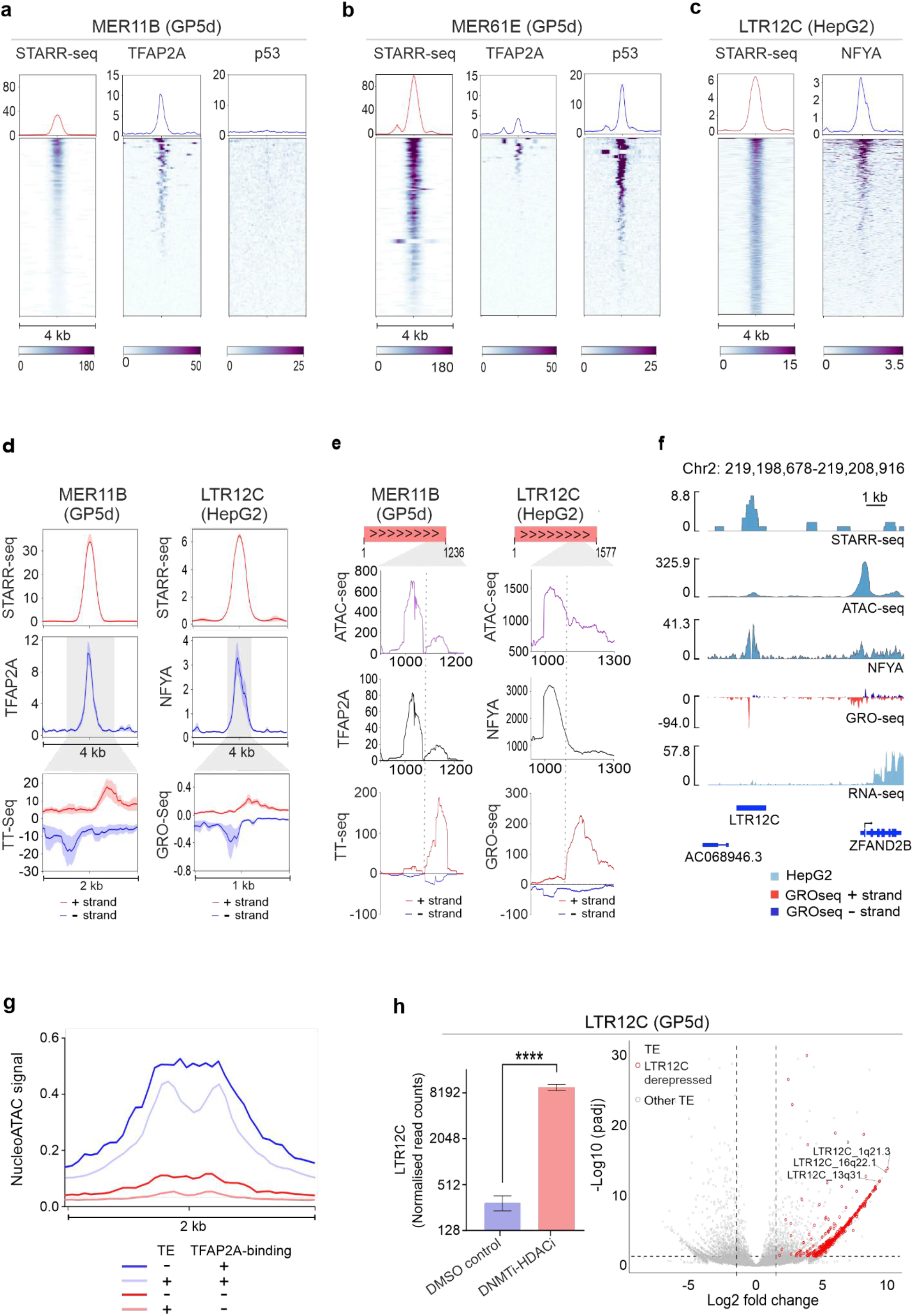
Distinct TFs regulate cell type-specific transcriptionally active TE enhancers. a) Heatmap of STARR-seq and ChIP-seq signals for TFAP2A and p53 at MER11B elements overlapping a STARR-seq peak in GP5d cells. Signal was plotted in ±2 kb flanks from the center of peaks. b) Heatmap for MER61E elements overlapping a STARR-seq peak in GP5d cells as in panel a. c) Heatmap of STARR-seq and ChIP-seq signals for NFYA at LTR12C elements overlapping a STARR-seq peak in HepG2 cells. Signal was plotted in ±2 kb flanks from the center of peaks. d) Left panel: metaplots of STARR-seq, TFAP2A ChIP-seq and TT-seq signals at MER11B elements overlapping a STARR-seq peak in GP5d cells. STARR-seq and TFAP2A ChIP-seq are plotted in a ±2kb region and TT-seq in a ±1kb region from the center of the STARR-seq peaks. Right panel: metaplots of STARR-seq, NFYA ChIP seq and GRO-seq signals at LTR12C elements overlapping a STARR-seq peak in HepG2 cells. STARR-seq and TFAP2A ChIP-seq are plotted in a ±2kb region and GRO-seq in a ±0.5kb region from the center of the STARR-seq peaks. All metaplots show the average signal with standard error. e) ATAC-seq, TFAP2A ChIP-seq and TT-seq reads in GP5d mapped to MER11B elements overlapping STARR-seq peaks were extracted and mapped to the MER11B consensus sequence. Left panel shows metaplots of the signals in a region from 900 to 1,236 bp of the MER11B consensus sequence. Similarly, ATAC-seq, NFYA ChIP-seq and GRO-seq reads in HepG2 mapped to LTR12C elements overlapping a STARR-seq peak were extracted and mapped to the LTR12C consensus sequence. Right panel shows metaplots of the signals in a region from 900 to 1,300 bp of the LTR12C consensus sequence. f) Genome browser snapshot showing STARR-seq, ATAC-seq, NFYA ChIP-seq and GRO-seq signal at a LTR12C element overlapping a STARR-seq peak in HepG2 cells. g) Metaplot of average nucleosome occupancy at GP5d STARR-seq peaks classified into four groups based on TFAP2A binding and overlap with TEs in a ±1kb region from STARR-seq peak centers. Nucleosome occupancy was determined by using NucleoATAC (see Methods for details). h) Changes in TE expression in GP5d cells upon DNMT and HDAC co-inhibition on a subfamily and locus level (see Methods for details). Left panel shows the normalized RNA-seq read counts for the LTR12C subfamily with and without the inhibitor treatment. Each bar represents mean value from three replicates and error bars correspond to standard deviation (two-sided unpaired t-test, p-value = 6.22259e-5. Right panel shows a volcano plot of differentially expressed TEs after the co-inhibition. 498 upregulated LTR12C elements are highlighted in red.

To confirm that the TEs overlapping STARR-seq peaks are functionally active enhancers *in vivo*, we utilized the data for nascent transcriptional activity, TT-seq from GP5d cells[28] and GRO-seq from HepG2 cells[29]. We observed divergent transcript initiation in GP5d cells at loci where STARR-seq peaks overlap with MER11B elements bound by TFAP2A (**Fig. 4d**). Similarly, STARR-seq peaks in HepG2 cells overlapping LTR12C elements and bound by NFYA were transcriptionally active (**Fig. 4d**). Next, we correlated the nascent transcriptional activity to TFAP2A binding at MER11B TE-STARR-seq peaks representing different clusters from **Fig. 2a**. In agreement with the STARR-seq signal and TFAP2A occupancy, TT-seq signal was strong in Cluster 4, moderate in Cluster 5 and weak in Cluster 1 (**Extended Data Fig. 4c**). Similarly, TT-seq signal correlated strongly with p53 binding in three different clusters of MER61E-enriched TE-STARR-seq peaks (**Extended Data Fig. 4d**). These results indicate that enhancer activity measured from the episomal STARR-seq assay correlates well with *in vivo* TF occupancy and nascent transcriptional activity around these TE overlapping enhancers.

As most copies of TEs in the genome are truncated, it is difficult to distinguish transcriptional activity between autonomously expressed full length TEs and truncated TE fragments that can still function as enhancers. To avoid this bias, we filtered unique reads from ATAC-seq, ChIP-seq and TT-seq/GRO-seq that mapped to annotated LTR elements and re-aligned the filtered reads to LTR consensus sequences (**Extended Data Fig. 5a, b**). We observed that accessible chromatin regions with enrichment for TFAP2A binding at MER11B and for NFYA binding at LTR12C elements associated with signals of divergent nascent transcription typical for enhancer RNAs as seen from GP5d TT-seq data and HepG2 GRO-seq data, respectively (**Fig. 4e**), further supporting the role of these specific subfamilies as cell type-specific enhancers derived from repetitive elements. **Fig. 4f** shows a representative example of transcriptionally active NFYA-bound LTR12C element located upstream of the *ZFAND2B* gene. NFYA is known to bind to promoter-proximal regions and to recruit pre-initiation complex to TSS[30], and the active transcription of the *ZFAND2B* gene in HepG2 cells is confirmed from the RNA-seq data (**Fig. 4f**). In conclusion, the clear position-specific enrichment of NFYA with respect to the signal of divergent transcription from GRO-seq data in HepG2 cells indicates that these repeats are functional enhancers resulting in enhancer RNA production (**Fig. 4d-4f**).

To further characterize the properties of TE-overlapping and non-TE enhancers, we analyzed ATAC-seq data in GP5d[31] for nucleosomal occupancy. For this, we classified GP5d STARR-seq peaks into four groups based on TFAP2A binding and overlap with TEs and discovered that TFAP2A-bound STARR-seq peaks generally have higher nucleosome occupancy than peaks not bound by TFAP2A (**Fig. 4g**). Interestingly, TE-enhancers bound by TFAP2A showed highly phased nucleosome pattern, whereas the non-TE-overlapping TFAP2A-bound peaks had an unphased, random nucleosomal distribution (**Fig. 4g and Extended Data Fig. 6a**), validating earlier reported pioneer activity of TFAP2A[32, 33]. Since the LTR12C elements showed higher enhancer activity only in HepG2 cells (*c*.*f*. **Fig. 3a and 4d**), we postulated that they might be epigenetically repressed and thus inactive in GP5d cells. To test this, we exposed GP5d cells to small molecule inhibitors for two epigenetic modifier enzymes, DNA methyltransferase (DNMT) and histone deacetylase (HDAC) followed by RNA-seq to specifically measure their effect on TE expression. Interestingly, DNMT and HDAC inhibition significantly upregulated LTR12C expression in GP5d cells at subfamily level (**Fig. 4h**, *left panel*) as well as locus level (**Fig. 4h**, *right panel*). To study the potential gene regulatory effect of these LTR12C-derived enhancers, we analyzed the expression of genes in the vicinity of LTR12C elements (+/-50 kb) from the RNA-seq data upon DNMT and HDAC inhibition. Interestingly, 89 out of 264 LTR12C-adjacent genes were differentially expressed, 70 of which were upregulated (**Extended Data Fig. 6b**). These results indicate that epigenetic de-repression – such as hypomethylation that is a general feature of human cancers – can lead to specific activation of distinct TEs and possible transactivation of adjacent genes, demonstrating how cancer cells can utilize TEs as *de novo* enhancers resulting in altered transcriptome profiles in the affected cells.

### The effect of DNA methylation on TE enhancer activity

DNA methylation is an important mechanism that suppresses TE activity in the genome. To study the effect of CpG methylation on TE enhancer activity in an unbiased manner, we performed genomic STARR-seq with *in vitro*-methylated libraries in HepG2 cells and compared enhancer activity of TEs by using our previously published STARR-seq data from HepG2 cells [17]. In total, 6568 peaks were called for the methylated library. Interestingly, we found that DNA methylation significantly repressed specific TE subfamilies, such as L1PA3, THE1B and THE1C (**Fig. 5a-c**). Motif enrichment analysis within the active THE1B and THE1C elements revealed several TF motifs harboring CG dinucleotides, such as BANP/ZBTB33 and HOXC13 (**Fig. 5d**). Importantly, in the motif enrichment analysis the active THE1B and THE1C elements that overlap with STARR-seq peaks were compared to elements of the same subfamilies that do not overlap with a STARR-seq peak, suggesting that regulatory differences can exist between active and silent TEs even within TE subfamilies. Supporting this, we observed stronger ZBTB33 binding to THE1 elements with enhancer activity in HepG2 cells compared to non-STARR-seq overlapping THE1 elements (**Fig. 5e**). Some homeodomain TFs such as HOXB13 can bind to both methylated and unmethylated motifs with a higher affinity to methylated CG[34], but further studies have suggested more complex interaction between HOXB13 and methylated DNA[35]. Interestingly, in the episomal STARR-seq assay, the HOXC13 motifs with TCG residues were enriched among the enhancers whose activity decreased upon DNA methylation (**Fig. 5d**). Moreover, out of the two TFs that can bind the CGCG sequence enriched within the THE1 elements, ZBTB33 has been shown to bind methylated CG[36] and BANP to non-methylated CG[37], highlighting the importance of studying the effect of DNA methylation on TE activity in a locus and context-specific manner. Finally, to analyze the methylation of THE1 elements at endogenous loci we utilized long-read single-molecule nanopore sequencing data, showing that genomic regions corresponding to active THE1-enhancers in HepG2 cells were less frequently methylated in HepG2 cells compared to GP5d cells (**Fig. 5f**). Taken together, our results suggest that specific TE subfamilies are affected by DNA hypomethylation, and their enhancer activity is controlled by TFs with differential binding preferences for methylated DNA.

**Figure 5.**
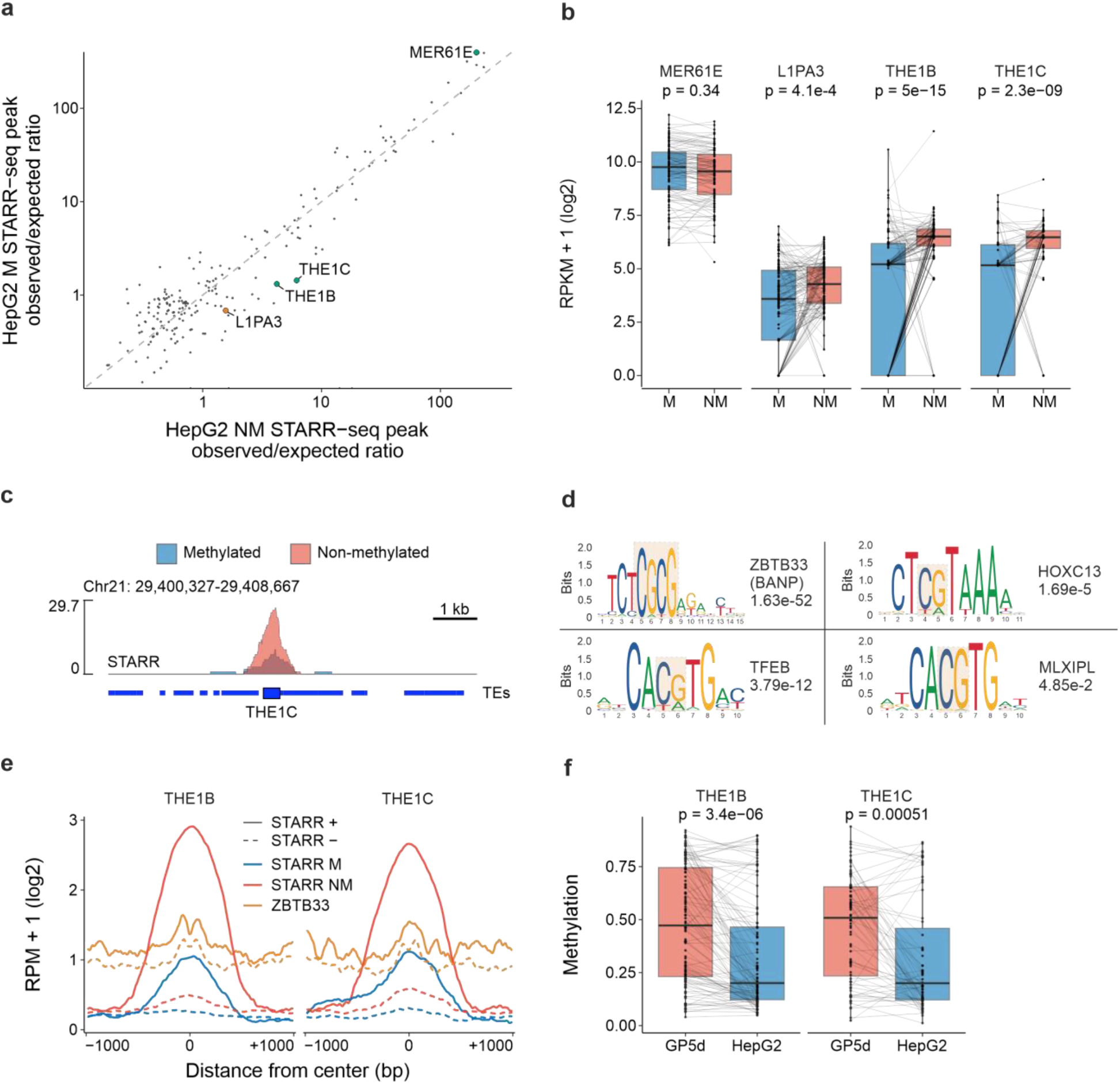
The effect of DNA methylation on TE enhancer activity. a) The effect of methylation on enhancer activity studied using non-methylated and *in vitro*-methylated genomic libraries in the STARR-seq assay in HepG2 cells. Observed/expected ratios plotted for STARR peaks overlapping with TE subfamilies, significantly different enrichment between non-methylated (NM) and methylated (M) libraries are labeled (BH-adjusted two-sided Fisher’s exact test FDR < 0.01). b) RPKM-transformed read counts from STARR-seq experiment in HepG2 cells using non-methylated (NM) and methylated (M) libraries at differentially enriched TE subfamilies from panel a. P-values were calculated with a two-sided, paired Wilcoxon test. The lower and upper hinges of the boxes represent the 25^th^ to 75^th^ percentiles, the midline is the median, and the whiskers extend from the hinges to the minimum and maximum values by 1.5 * IQR (interquartile range, i.e. distance between the 25^th^ and 75^th^ percentiles). c) Example genome browser snapshot of a THE1C element showing a reduction in STARR signal from a methylated genomic library compared to non-methylated library in HepG2 cells. d) Motif enrichment within THE1B and THE1C elements overlapping a STARR peak summit in HepG2 cells compared to non-STARR-overlapping THE1B and THE1C elements. As background sequences, 5000 THE1B and THE1C elements that did not overlap with a STARR-seq peak summit were randomly selected. The sequence logo with CG dinucleotides highlighted, name of the TF and the E-value from AME (see Methods for details). Only significant human-specific motifs are shown (E-value < 0.05). e) Metaplot of RPM-normalized signals for STARR-seq using non-methylated (STARR NM) and methylated (STARR M) libraries as well as ZBTB33 ChIP-seq at THE1B and THE1C elements in HepG2 cells. Coverage was calculated in a ±1kb region from the center of the elements, log_2_-transformed and a rolling mean was calculated with a 25bp window. Solid line shows the signal at the THE1B and THE1C elements overlapping a STARR peak summit, and dashed line shows the signal at 1000 randomly sampled THE1B and THE1C elements that do not overlap with a STARR peak summit. f) Boxplots of CpG methylation of THE1B and THE1C elements in GP5d and HepG2 cells analyzed from NaNOMe-seq data. Smoothed methylation signal for GP5d and HepG2 was taken from all the elements with a STARR-seq peak summit overlap in HepG2 and compared. Boxplot features are as in panel **b**. P-values are from a two-sided paired Wilcoxon test.

### *In silico* prediction of TE enhancer–gene contacts show widespread changes in gene expression

To analyze the effect of TE enhancers on gene expression, we utilized the activity-by-contact (ABC) model[38] for *in silico* prediction of genomic contacts between identified TE enhancers and potential target genes (see Methods). In total, the model predicted 29,967 contacts in GP5d cells, 1,206 of which overlapped with a STARR-seq peak summit and were selected for further analysis. Of these, 486 overlapped with a TE. As expected, most of the predicted contacts mapped to Cluster 1 (*c*.*f*. **Fig. 2a**) with almost half of the STARR-seq peaks in Cluster 1 showing at least one predicted enhancer–target gene contact. This is consistent with the active epigenetic signals (ATAC+, H3K27ac+) enriched in Cluster 1 which are also used by the ABC model for contact prediction (**Extended Data Fig. 7a**), speaking for the activity of these TE elements in GP5d cells.

Since the MER11 elements were strongly overrepresented and active in several clusters in GP5d cells (*c*.*f*. **Fig. 2c, 3a, 4a, 4d**), we analyzed if the genes predicted to be connected to these elements show differential expression between GP5d and HepG2 cells. We found that the majority of the predicted MER11 target genes were overexpressed in GP5d cells in comparison to HepG2 (**Fig. 6a**). The expression of MER11-associated genes was also higher in GP5d colon cancer cells compared to normal colon epithelium HCoEpiC cells (**Extended Data Fig. 7b**), supporting the observation that MER11 subfamilies can function as active enhancers in GP5d cells. For example, a MER11B-element in chromosome 3 with a strong STARR-seq signal, active enhancer marks and TFAP2A binding in GP5d cells has a predicted association to several genes that are strongly expressed in GP5d cells (**Fig. 6c**). In HepG2 cells, however, the MER11B-enhancer is not active, and the expression of these genes is low despite the active epigenetic marks at their promoters (**Fig. 6c**).

**Figure 6.**
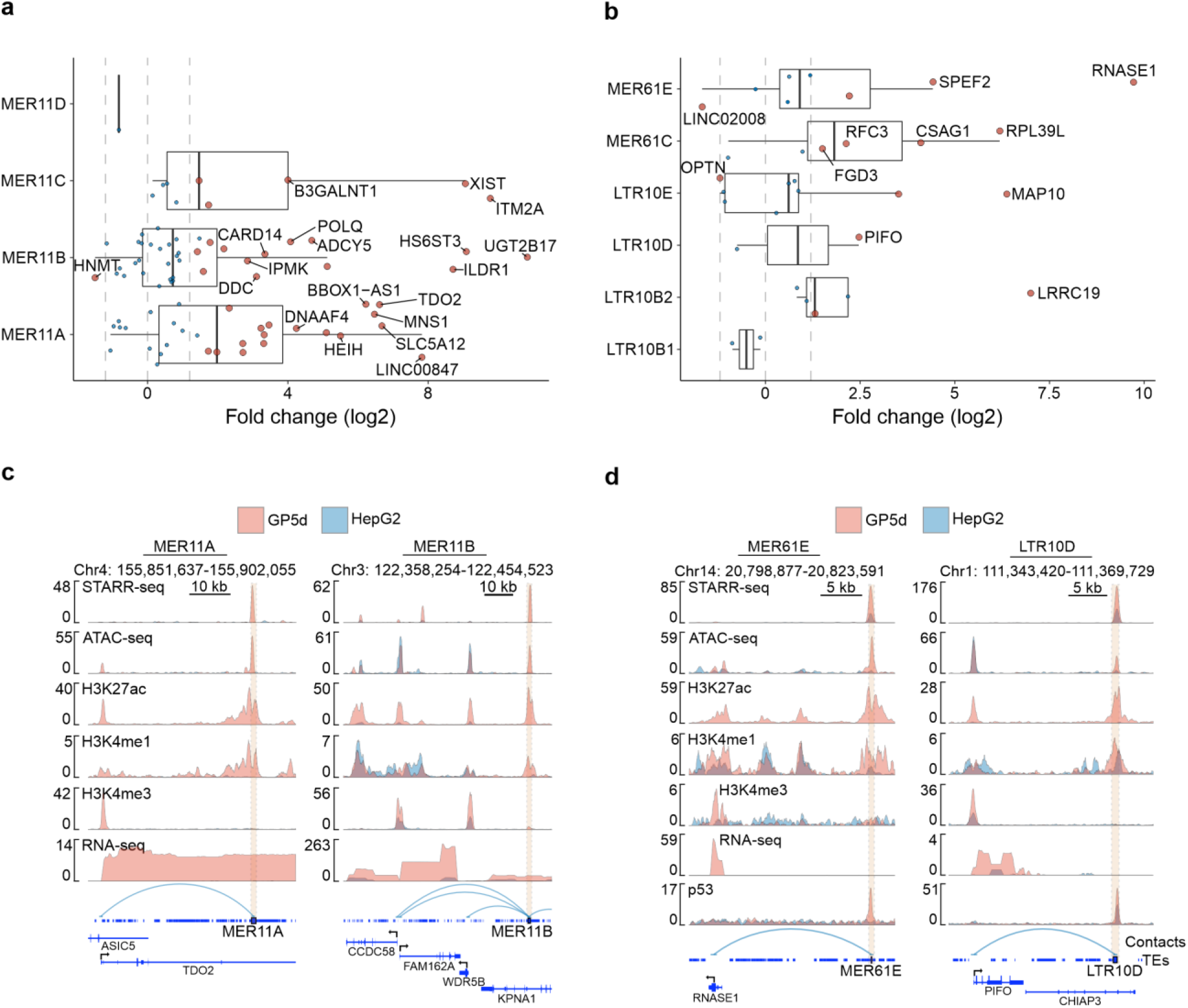
*In silico* prediction of TE enhancer–gene contacts show widespread changes in gene expression. a) Differential expression of genes with predicted contacts to MER11 subfamily TEs. Log_2_ fold change shown from RNA-seq data for GP5d vs. HepG2 cells. Red points mark the significantly differentially expressed (|Log_2_ fold change| > 1.2 and BH-adjusted FDR < 0.05), blue points are insignificant. Log_2_ fold change of ±1.2 and 0 is marked with a dashed line. The lower and upper hinges of the boxes represent the 25^th^ to 75^th^ percentiles, the midline is the median, and the whiskers extend from the hinges to the minimum and maximum values by 1.5 * IQR (interquartile range, i.e. distance between the 25^th^ and 75^th^ percentiles). b) Differential expression of genes with predicted contacts to p53-specific TEs from RNA-seq data between GP5d WT vs. HepG2 cell lines. Plot elements are similar to panel a. c) Genome browser view of an active MER11B element with high STARR-seq signal and 21 predicted contacts to genes in GP5d cells (all contacts not shown). ATAC-seq tracks for chromatin accessibility and ChIP-seq for histone marks and TFAP2A binding, RNA-seq data showing the expression of four genes that are predicted to be regulated by the MER11B element are shown in GP5d and HepG2 cells. Of note, HepG2 lacks the active signals at the MER11B-element and the genes show low expression in HepG2 despite the epigenetic marks of active promoters (ATAC, H3K4me3). d) Genome browser view of one of the overexpressed genes in GP5d WT vs. GP5d p53-null cells, *PIFO*, with a predicted contact with a p53-specific LTR10D element. The lower STARR-seq and p53 ChIP-seq signals in HepG2 vs. GP5d cells correlate with lower expression of the *PIFO* gene.

As TEs enriched within many STARR-seq peak clusters are highly p53-specific, such as MER61 and LTR10 subfamilies, we studied the expression of genes with a predicted contact to these TEs. For this, we compared gene expression between wild-type GP5d and HepG2 cells and observed that the genes with predicted association to MER61 and LTR10 subfamilies also showed upregulation in GP5d cells (**Fig. 6b**), consistent with the higher enrichment of active enhancer features such as ATAC-seq signal and ChIP-seq signals for H3K27ac and H3K4me1 observed in GP5d cells compared to HepG2 cells at some of these elements (**Fig. 6d**). Previously, it has been shown that despite the presence of strong p53REs in TEs, p53 binding rarely resulted in transactivation of genes[39]. We also show that relatively few of the highly enriched p53s-specific TEs had a predicted contact to a gene, but some of them were associated with an overexpressed gene and active enhancer marks in GP5d cells (**Fig. 6b, d**). When comparing the expression of the genes with predicted contact to a p53-associated TE between wild type and p53-null GP5d cells, only few of the genes showed significant difference in expression (**Extended fig. 7c**). Primary cilia formation gene, *PIFO*, had a predicted contact to a p53RE-containing LTR10D element and showed a significantly altered expression after p53 depletion as well as in GP5d cells compared to HepG2 (**Fig. 6b, d; Extended fig. 7c**). In GP5d, the LTR10D element had an active enhancer profile, whereas in HepG2 cells the H3K27ac signal was lacking, suggesting a poised enhancer state[40] and consequently lower expression of the gene (**Fig. 6d**). This suggests that p53RE-containing TEs may also have a minor role in regulating gene expression as a response to p53. In conclusion, our results suggest that the active TEs may cause widespread changes in gene expression in cancer cells.

## Discussion

It is well-established that TEs are a rich source of gene regulatory elements[2, 27] and have contributed to the evolution of gene regulatory networks[13]. However, the role and the extent of TEs as onco-exapted enhancers is less well understood. Here, we studied functional enhancer activity of TEs in two endodermal origin cancers by combining data from a quantitative whole genome enhancer assay to extensive epigenetic analyses. The advantage of the episomal STARR-seq assay is that it enables measuring the regulatory potential of each genomic element in an unbiased manner, allowing the functional analysis of endogenously silent repetitive elements, whereas the epigenome analysis reveals the regulatory status of each element in the endogenous genomic context. We identified specific TE subfamilies that are enriched in colon and liver cancers with a notable specificity in active TE enhancers between these two cancer types, suggesting that the TE enhancers can be activated in a cell type-specific manner.

TEs are under strict epigenetic control in normal somatic cells through DNA methylation, repressive histone modifications and RNA-mediated silencing, mediated through TFs such as p53 and KRAB-ZNFs[21, 22]. Two opposing hypotheses have been proposed for TE activity and exaptation based on the evolutionary age of the elements and their epigenetic status[41]: exaptation hypothesis predicts that more conserved and older TEs are more enriched for active histone marks due to their exaptation to regulatory roles during genome evolution, and defense hypothesis predicts that young TEs will be more enriched for repressive marks due to their potential disruptive activity. This classification was based on histone modifications only, whereas here we have also used nascent transcription data and unbiased enhancer activity measurements for detailed characterization of TE activity. In favor of the defense hypothesis, we observed strong STARR-seq activity, more histone modifications, and nascent transcription at relatively young LTR subfamilies as opposed to older TE subfamilies. However, our results highlight the context-specificity of TE activation, since different copies of the same subfamily can be associated to either active or repressed chromatin state as revealed by our detailed clustering analysis. Importantly, during tumorigenesis, TEs can escape their silencing for example in response to epigenetic dysregulation, providing scope for onco-exaptation of previously suppressed TEs. This emphasizes the need for detailed understanding of their suppressing and activating mechanisms in different cancer types, as done here for colon and liver cancers.

Majority of the enriched TE enhancers in both colon and liver cancer cell lines represent the ERV1 family of LTRs with significantly more STARR-seq peaks overlapping with LTR elements than expected from a random distribution. This is consistent with earlier reports demonstrating that LTR elements contribute to ∼39% of TF binding sites in both human and mouse genomes despite comprising only 8% of the human genome[27]. Transcripts originating from ERV1 elements have been detected in several cancer types, and high level of ERV1 expression was associated to poor outcome in kidney cancer[42, 43]. The widespread ERV expression detected in human cancers speaks for the global de-repression of epigenetic silencing of these elements. Interestingly, nascent transcription from enhancers has been shown to be more predictive of enhancer activity than enrichment of active histone marks[44]. Here, we showed that enhancer activity of TEs measured using the episomal STARR-seq assay correlates well with nascent transcription and active histone marks from the respective genomic loci. Our data suggests that the same transcriptionally active TEs that evade epigenetic suppression can also be functionally active enhancers, warranting further studies about their significance in tumorigenic processes.

The active enhancers detected from GP5d colon cancer cells and HepG2 liver cancer cells using STARR-seq were enriched for distinct as well as common TE subfamilies. The common elements, such as primate-specific MER61 and LTR10 subfamilies, were strongly enriched for p53 motif. p53 is functionally active in both cell lines, and the common active TEs overlapped with p53 binding in the genome. The common elements were among the most highly enriched subfamilies in both cell lines and had the highest STARR-seq activity but only weak enrichment for the active histone marks, consistent with earlier reports from human and mouse cells[17, 45]. The enrichment of TE subfamilies correlated with the proportion of their genomic copies harboring p53 binding motifs. Previous studies have shown that active and functional p53 enhancers are characterized by a single canonical p53 motif without binding of coregulatory TFs[46], commensurate with our findings in Clusters 4 and 5 with strong enrichment of p53 motifs. A recent study found that p53 enhancers are also likely to be independent of interaction via Mediator[47], further supporting independent function of p53 as a transcriptional regulator.

In the context of TE regulation, both silencing and activating roles have been suggested for p53[24, 48]. p53 is known to be a pioneer factor[49, 50] that can preferentially bind to regions of high nucleosome occupancy[51] and closed chromatin enhancers[17]. It has been shown previously that TE-associated p53-occupied p53REs were stronger than non-TE associated p53REs, especially at LTR10B and MER61 elements, but p53 binding at TEs did not cause changes in expression of target genes[39]. Our findings were in line with this, showing that the expression of *in silico*-predicted target genes of MER61 and LTR10 enhancers was largely unaffected by p53 depletion. In our study, strong p53REs were found within enhancers and repressed chromatin, possibly explaining the strong STARR signal, high H3K9me3 and low chromatin accessibility in Cluster 4. In conclusion, our results support both silencing and activating functions for p53, showing that majority of the p53-enriched TEs that show enhancer activity in the episomal STARR-seq assay are associated with repressive histone marks at the endogenous chromatin loci, but that specific upregulation of TE-associated predicted p53 target gene, such as PIFO, was also detected.

Our analysis of the STARR-seq peaks also revealed active TE enhancers specific to each cancer type. MER11 subfamilies of the LTR class (MER11A, B, C and D) were specifically active in GP5d colon cancer cells showing enrichment for TFAP2A motif and overlap with TFAP2A binding in the genome. Nascent transcriptional activity from the MER11B elements correlated with TFAP2A ChIP-seq and STARR-seq signals, further speaking for the enhancer activity of MER11 elements. Interestingly, CRISPR screen data from Cancer Dependency Map (DepMap) collection shows that TFAP2A depletion has the strongest growth inhibitory effect in GP5d cells across 55 colorectal cell lines (see **Extended Data Fig. 6c**) (CRISPR Chronos score = - 0.2586)[52], suggesting that TFAP2A-controlled transcriptional programs are critical for GP5d proliferation. Previous studies have reported pioneering activity for TFAP2A[32, 33] demonstrating TFAP2A and its isoforms as strong nucleosomal binders[33]. However, the strong nucleosomal phasing that we observed specifically at the TFAP2A-bound TE loci is a novel finding, and the mechanism of the TE-specific nucleosome phasing in the context of TFAP2A pioneer activity warrants further studies.

In HepG2 liver cancer cells, we found that the LTR12 subfamilies were specifically enriched within the STARR-seq peaks. Consistent with a previous report, LTR12 subfamilies were highly enriched for NFY family motifs[53] and our data shows nascent transcriptional activity from NFYA-bound LTR12-enhancers. NFYA is essential for maintaining nucleosome-depleted regions at promoters of NFYA target genes and preventing sub-optimal transcription initiation from alternative TSSs by directing transcriptional machinery to the correct TSS[30]. The role of LTR12C as an alternative promoter has also been previously studied[53, 54]. Interestingly, we show that NFYA was associated to enhancer activity from the STARR-seq-identified LTR12 elements with high enrichment of the LTR12C within the STARR-seq peaks, raising the question whether LTR12C elements can also function as promoter-enhancers that were reported earlier[55].

The general mechanisms for TE silencing in somatic cells include DNA methylation, histone modifications, and RNA-mediated silencing. Interestingly, our results suggest that tissue-specific TE activation is associated to specific de-repression of epigenetic silencing. For example, LTR12C elements were found to be more active in HepG2 compared to GP5d cells, but inhibition of epigenetic regulatory enzymes DNMT and HDAC resulted in strong and specific upregulation of LTR12 elements in GP5d cells. Importantly, this also led to increased expression of endogenous genes in the vicinity of LTR12 elements, suggesting that cancer cells can utilize TEs for controlling the expression of endogenous genes. Furthermore, differential CpG methylation was observed between HepG2 and GP5d cells at THE1 elements that were shown to be more active in non-methylated conditions compared to methylated DNA in HepG2 cells. Taken together, our results demonstrate how re-activation of specific TE elements can occur through destabilization of epigenetic repressive mechanisms.

Due to the inherent difficulties of mapping the short-read sequences to the repetitive sequences within TEs, the results in this study may underestimate the actual extent of functional repertoire of TEs. Here, we only retained uniquely mapping reads, but methods to improve read assignment to TEs[56, 57] and peak calling for STARR-seq have been developed[58]. However, despite the lower mappability of evolutionarily young subfamilies, we found that relatively young LTR subfamilies were highly enriched in the STARR-seq data. Improvements in both short- and long-read sequencing technologies will benefit the future studies of TEs, reducing the limitations inherent to short-read sequencing methods[56, 57] and peak calling for STARR-seq have been developed[58]. However, despite the lower mappability of evolutionarily young subfamilies, we found that relatively young LTR subfamilies were highly enriched in the STARR-seq data. Improvements in both short- and long-read sequencing technologies will benefit the future studies of TEs, reducing the limitations inherent to short-read sequencing methods.

In conclusion, we studied the extent of TE enrichment in a genome-wide, unbiased functional enhancer assay together with orthogonal functional genomics methods for chromatin accessibility, histone modifications and transcription factor binding. We found that specific TE subfamilies are highly overrepresented among the active enhancers, showing remarkable tissue-specificity that is controlled by distinct TFs. The contribution of these TE-derived enhancers to tumorigenic processes warrants further studies, but our results provide first evidence of widespread exaptation of TEs for cancer type-specific functional enhancer activity.

## Materials and Methods

### Data acquisition

GP5d and HepG2 STARR-seq data was acquired from under GEO accession GSE180158[17]. For GP5d, genomic and p53-null STARR-seq data were downloaded (GSM5454433 and GSM5454435) and for HepG2 genomic STARR replicates 1 and 2 were downloaded (GSM5454437 and GSM5454438). GP5d ChIP-seq data for H3K27ac (GSM5454417), H3K9me3 (GSM5454420), H3K27me3 (GSM5454428), 5-FU treated mIgG (GSM5454414), untreated p53 (GSM5454412) and 5-FU treated p53 (GSM5454413) were acquired from the same study. GP5d H3K4me1 (GSM1240814) was obtained with the GEO accession GSE51234[59].

HepG2 ChIP-seq data for H3K27ac, H3K27me3, H3K36me3, H3K4me1, H3K4me3, H3K9me3, p53, NFYA and ZBTB33 replicate 1 and 2 were downloaded from ENCODE[60] (https://www.encodeproject.org/) with fastq file accessions ENCFF000BFD, ENCFF001FLQ, ENCFF001FMA, ENCFF000BEX, ENCFF901NZE, ENCFF000BFK, ENCFF257UIJ, ENCFF081VHA, ENCFF204TCE and ENCFF745CTV respectively. Replicate 1 for HepG2 ATAC-seq was downloaded with accessions ENCFF664UPL and ENCFF289UIB for read file 1 and 2 respectively. Three replicates for HepG2 RNA-seq (ERR6351780, ERR6351781 and ERR6351782) were downloaded under ENA accession PRJEB31262[61].

NCBI genome annotation files for GRCh38 were downloaded from Illumina iGenomes (http://igenomes.illumina.com.s3-website-us-east-1.amazonaws.com/Homo_sapiens/NCBI/GRCh38/Homo_sapiens_NCBI_GRCh38.tar.gz).

A gene annotation GTF file was acquired from Gencode Release 36 for the reference chromosomes (https://ftp.ebi.ac.uk/pub/databases/gencode/Gencode_human/release_36/gencode.v36.annotation.gtf.gz). The GTF file was transferred into a BED file with gtfToBed.sh (github.com/timplab/nanoNOMe/blob/isac/analysis/annotations/gtfToBed.sh) and a TSS and a gene body BED files were created with a script adapted from https://github.com/isaclee/nanoNOMe/blob/master/snakemake/downloaded_data_parse.smk[6 2].

A repeatMasker.txt (2021-09-03) file was downloaded from the UCSC table browser (https://genome.ucsc.edu/cgi-bin/hgTables). Only transposable element-derived repeat classes (LINE, SINE, LTR, and DNA) were retained and a file in BED format was created from the table, totaling 4745258 annotated repeats[63]. MER11B and LTR12C consensus sequences were acquired from RepBase[64] (https://www.girinst.org/repbase/update/index.html).

GRCh38 chromosome sizes file (2020-03-13) file was downloaded from UCSC (https://hgdownload-test.gi.ucsc.edu/goldenPath/hg38/bigZips/latest/).

Unified GRCh38 blacklist BED file (ENCFF356LFX, release 2020-05-05) was downloaded from ENCODE project (https://www.encodeproject.org/).

A genome index was created with bowtie2-build, with chr1-22, X, Y and M fasta files. Alternative, unlocalized and unplaced alternative loci scaffolds were discarded in indexing.

Transcription factor motifs were acquired from JASPAR 2022 CORE non-redundant vertebrate annotations (https://jaspar.genereg.net/download/data/2022/CORE). The position weight matrices in MEME format were used for downstream motif analyses.

Predicted LTR p53 binding site percentages were downloaded from[24]. TE age estimation data were downloaded from the TEanalysis pipeline[65], (https://github.com/4ureliek/TEanalysis/blob/master/Data/20141105_hg38_TEage_with-nonTE.txt).

### Cell culture

GP5d cells (Sigma, 95090715) were cultured in DMEM (Gibco, 11960-085) supplemented with 10% FBS, 2 mM L-glutamine (Gibco, 25030024) and 1% penicillin-streptomycin (Gibco, 15140122). HepG2 (ATCC, HB-8065) were cultured in MEM (Gibco, 11544456) supplemented with 10% FBS, 2 mM L-glutamine and 1% penicillin-streptomycin. HCoEpiC (ScienCell, 2950) were cultured in Colonic Epithelial Cell Medium (ScienCell, 2951) as per vendor’s guidelines.

### Genomic STARR-seq

Genomic STARR-seq experiment was performed in HepG2 cells as previously described[17] by using methylated input DNA library (two replicates, 170 million cells per replicate). One μg of methylated input library DNA was transfected per million HepG2 cells. In brief, 10 million HepG2 cells were seeded per 15-cm plate in media without antibiotic a day before transfection. Methylated plasmid DNA was mixed with Transfectin (Bio-Rad, 170-3351) at a 1:3 ratio in Opti-MEM medium (Gibco, 11524456), incubated for 15 minutes at room temperature and added dropwise to the cells. Cells were harvested after 24 hours after transfection and total RNA isolated using the RNeasy Maxi kit (Qiagen, 75162) with on-column DNase I digestion. Dynabeads mRNA DIRECT Purification kit (Invitrogen, 61012) was used to purify poly(A)+ RNA. poly(A)+ RNA was treated with DNase by using TurboDNase (Ambion, AM2238) followed by purification by using RNeasy Minelute kit (Qiagen, 74204). STARR-seq reporter library was prepared by following protocol as described in ref.[17] and paired end sequenced.

### ChIP-seq, ATAC-seq, NaNoME-seq and Hi-ChIP

ChIP-seq was performed as previously described[17] by using the following antibodies (2 μg per reaction) for: H3K4me3 (Sigma-Aldrich, 07-473), H3K36me3 (Diagenode, C1541092-10), TFAP2A (Abcam, AB52222), mouse IgG (Santa Cruz Biotechnology, sc-2027) and rabbit IgG (Santa Cruz Biotechnology, sc-2025). In brief, GP5d cells were formaldehyde cross-linked for 10 minutes at room temperature. Sonicated chromatin was centrifuged, and the supernatant was used to immunoprecipitated DNA using Dynal-bead coupled antibodies. Immunoprecipitated DNA was purified and used for ChIP-seq library for Illumina sequencing. The libraries were single-read sequenced on NovaSeq6000.

ATAC-seq library was prepared by using 50000 GP5d cells by using protocol described earlier[66]. GP5d cells were washed with ice-cold PBS and cells were resuspended in 50 μl lysis buffer. Cells were incubated for 10 minutes on ice. The pellet was resuspended in 2x tagmentation buffer (Illumina kit) and incubated at 37°C for 30 minutes. DNA was purified by using MinElute purification kit and eluted in nuclease free water. Optimal number of amplification cycles was determined by qPCR. Samples were amplified by using Nextera library preparation kit (Illumina) and sequenced paired end.

NaNoME-seq was performed to profile CpG methylation and chromatin accessibility (GpC methylation) in GP5d cells as described earlier[61]. GP5d cell nuclei were isolated and treated with GC methylase M.CviPI (New England Biolabs, M0227) as described[61]. Following GC methylation, DNA was isolated from nuclei by using phenol chloroform extraction protocol and sequencing library was prepared using the 1D genomic DNA by ligation kit (SQKLSK109) according to manufacturer’s protocol and 50 fmol of adaptor-ligated genomic DNA was loaded to the flow cell for sequencing.

H3K27ac HiChIP was performed as previously described in ref.[67]. In brief, GP5d cells were formaldehyde cross-linked for 10 minutes at room temperature and nuclei were isolated from crosslinked cells by 30 minutes of lysis at 4 °C. Isolated nuclei from 15 million cells were permeabilized in 0.5% SDS for 10 minutes at 62 °C and SDS was quenched by using Triton X-100 for 15 minutes at 37 °C. Chromatin was digested with MboI restriction enzyme (New England Biolabs, R0147) at 37 °C for 2 hours and then heat-inactivated at 62 °C for 20 minutes. Chromatin was incubated with a DNA ligase (New England Biolabs, M0202) for 4 hours at room temperature for proximity ligated contact formation followed by centrifugation. Proximity ligated chromatin pellet was sonicated and immunoprecipitated using 5 μg of H3K27ac antibodies (Diagenode, C15410196). Immunoprecipitated fragments were Biotin labeled (Invitrogen, 19524016) and captured by Dynabeads MyOne Streptavidin C1 beads (Invitrogen, 65001). Immunoprecipitated DNA was adaptor-labeled by using Tn5 transposase (Illumina) and PCR amplified. The libraries were paired-end sequenced on NovaSeq 6000.

### DNMT and HDAC inhibitor treatment and RNA-seq

DNMT and HDAC inhibition in GP5d cells were performed as described earlier[53]. GP5d cells were seeded in 6 well plates and treated with 500 nM/L 5-aza-2’-deoxycytidine (MedChemExpress, HY-A0004) or DMSO (Fisher, BP231). 5-aza-2’-deoxycytidine containing media were replaced each day for three days. SB939 was added at a concentration of 500 nmol/L (MedChemExpress, HY-13322) and cell were harvested after 18 hours. Total RNA was extracted using RNeasy Mini kit (Qiagen) according to the manufacturer’s instructions in three biologicals replicates. RNA-seq libraries were prepared using 1 μg of total RNA input using KAPA stranded mRNA-seq kit for Illumina (Roche) as per manufacturer’s instruction and paired-end sequenced on NovaSeq 6000 (Illumina).

### Genomic STARR-seq analysis

FastQC v.0.11.9 was used for quality control and determination of read lengths of raw data[68]. For GP5d, the input plasmid control reads were trimmed with Trimmomatic v.0.39 from ∼76bp to 36bp, matching the read length of reporter cDNA to avoid biases in mappability[69]. For HepG2, the two technical replicates were combined before alignment. Bowtie2 v.2.4.1 was used to map the paired-end STARR-seq reads to the reference human genome (hg38/GRCh38) with the (bowtie2 --maxins 1000)[70]. Duplicates were marked with Picard v.2.23.4 (MarkDuplicates - REMOVE_DUPLICATES false -ASSUME_SORTED true) and quality metrics were determined with picard CollectMultipleMetrics (http://broadinstitute.github.io/picard/). Samtools v.1.7 was used to filter non-concordant reads and reads with a MAPQ smaller than 20 (samtools view -h - F 1024 -q 20)[71]. MACS2 v2.2.7.1 was used for calling peaks with options –f BAMPE –g hs and the STARR input as a control[72]. Peaks overlapping with ENCODE blacklisted regions were removed from the narrowPeak and summit files with bedtools v.2.29.2 (bedtools subtract -A)[73]. Genome read coverage was calculated with bedtools v.2.29.2 (bedtools coverage –pc -bg) and bedGraphToBigWig v.377 was used to create a bigwig file[74]. A RPKM normalized bigwig file was created with deepTools v.3.5.0 (bamCoverage --binSize 50 --normalizeUsing RPKM -- effectiveGenomeSize 2913022398)[75]. Final mapped read statistics are listed in **Supplementary table 5**.

### ATAC-seq analysis

The raw reads were mapped with bowtie2 v.2.4.1 (bowtie2 --very-sensitive) to the reference human genome (hg38/GRCh38). Reads mapped to the mitochondrial genome were removed with removeChrom.py script (https://github.com/jsh58/harvard/blob/master/removeChrom.py). Picard v.2.23.4 was used to remove duplicates (MarkDuplicates -REMOVE_DUPLICATES false -ASSUME_SORT_ORDER coordinate) and analyzing insert sizes with CollectInsertSizeMetrics. Samtools v.1.7 was used to filter reads with MAPQ smaller than 10 and remove marked duplicates (samtools view -F 1024 -b -q 10). Peaks were called with MACS2 v.2.2.7.1 (macs2 callpeak -f BAMPE -g hs --keep-dup all). Removal of blacklisted regions, coverage calculation, conversion to bigwig and normalized coverage file creation was performed as in STARR-seq data processing. Final mapped read statistics are listed in **Supplementary table 5**.

### ChIP-seq analysis

ChIP-seq data processing was performed as in ATAC-seq except for removing the reads mapping to the mitochondrial genome and read filtering, where reads with a MAPQ smaller than 20 were removed. Final mapped read statistics are listed in **Supplementary table 5**.

### Nanopore-seq analysis

The nanopore data was processed with ONT_hg_pipe v.0.1.0 (Palin 2018, unpublished, https://github.com/kpalin/). The GP5d nanopore data was basecalled with Guppy 5.0.17 with the super-accurate basecalling model and a minimum q-score of 10. The reference genome was indexed and the basecalled reads were aligned to the reference genome with minimap2 2.16 (minimap2 –x map-ont)[76]. Quality controls were performed with nanoplot v.1.20.0[77] and Samtools 1.9[71]. After alignment, methylation was called with nanopolish v.0.11.1[78]. The cpggpc_new_train branch in GitHub (https://github.com/jts/nanopolish/tree/cpggpc_new_train) was used to call both CpG and GpC methylation (nanopolish call-methylation -q cpggpc). The resulting table was processed to a BED format (mtsv2bedGraph.py -q cpggpc --nome) and to methylation frequency table formats for CpG and GpC methylation (parseMethylbed.py frequency -v -m cpg and parseMethylbed.py frequency -v -m gpc), using previously published scripts[62]. The resulting methylation tables were converted to bedGraph and bigwig formats with a custom script (mfreq_to_bw.R), utilizing bedGraphToBigWig v.377[74]. The CpG methylation frequency tables were loaded into R and smoothed with bsseq v.1.28.0 (BSmooth ns = 50, h = 1000, maxGap = 100000)[79]. The GpC methylation calling was performed for another project and was not used in this study.

### RNA-seq analysis

FastQC v.0.11.9 was used for quality control. Pseudoalignment and counting was performed with Salmon v.1.8.0[80]. Reads were also aligned with STAR v.2.5.3a[81] by using the SQuIRE pipeline v. 0.9.9.92[82]. Read counts were calculated with featureCounts v.2.0.1[83]. Differential expression analysis was performed with DESeq2 v.1.32.0[84].

### Hi-ChIP-seq analysis

Hi-ChIP data for GP5d H3K27ac was processed with HiC-Pro v3.1.0[85] with default parameters and hg38 MboI restriction sites .bed file as input. allValidPairs output from HiC-Pro was converted to a .hic file with the hicpro2juicebox.sh script from HiC-Pro, with juicer tools v.1.22[86]. KR-normalized matrices were extracted from the .hic file with juicebox_dump.py from ABC v.0.2.2[38] and powerlaw fit was calculated with the compute_powerlaw_fit_from_hic.py script.

### GP5d TT-seq and HepG2 GRO-seq

GP5d TT-seq data (GSM4610669) was acquired under GEO accession GSE152291[28]. HepG2 GRO-seq raw data were downloaded under GEO accession GSM2428726 (SRR5109940)[29]. GRO-seq reads were trimmed to remove A-stretches originating from the library preparation by using the Trim Galore v.0.6.7. Sequence reads shorter than 25 bp and quality score less than 10 were discarded. GRO-seq reads were aligned to the hg38 genome assembly using bowtie2 v.2.2.5 and strand specific bigwig files were created with Samtools v.1.9.

### TEtranscripts and Telescope RNA-seq analysis

FastQC v.0.11.9 was used for quality control. Reads were also aligned with STAR v.2.5.3a[81] by using the SQuIRE pipeline v. 0.9.9.92[82] TEtranscripts v.2.2.1 was run on the SQuIRE alignment output with the following flags: –mode multi –stranded reverse. The log2 fold change values output by DESeq2 v1.32.0 were used for TE subfamily expression analysis. Telescope v.1.0.3.1 analysis was performed on SQuIRE alignment output. The “telescope assign” command was used to quantify TE expression. The log2 fold change values output by DESeq2 v1.32.0 were used in subsequent analysis. Nearby genes were associated to TEs with GREAT v.4.0.4[87].

### TE enrichment analysis

STARR-seq peak summits for GP5d and HepG2 were overlapped with repeatMasker annotations with GenomicRanges R package v.1.44.0[88] and the count of overlaps for each subfamily of TEs was summed. Only TE subfamilies with 5 or more overlaps in total were retained for further analysis.

The expected frequency of overlaps was calculated by shuffling the STARR-seq peak summits 1000 times with R package bedtoolsr v.2.29.0-3[89], excluding masked regions in from the BSgenome.Hsapiens.UCSC.hg38.masked v.1.3.993 package and keeping the shuffled features in the same chromosomes (-chrom). The mean frequency of shuffled overlaps for each TE subfamily was calculated, giving the expected frequency of overlaps. The observed/expected ratio was calculated by dividing the overlap of STARR-seq peaks by the expected overlaps for each subfamily.

Statistical significance for each TE subfamily enrichment was calculated with a one-sided binomial test with binom_test from the rstatix package v.0.7.0[90]. The number of overlaps of STARR-seq peaks was set as the number of successes, the count of all STARR-seq peaks as the number of trials and the fraction of shuffled overlaps for a subfamily from all the shuffled overlaps was the probability of success. P-values were adjusted with Benjamini-Hochberg correction[91]. Significance was determined as FDR < 0.01.

The differential enrichment for TE subfamilies was calculated with a two-sided Fisher’s exact test with the count of overlaps for a TE subfamily as the observed and the sum of 1000 shuffled overlaps as the expected frequency. P-values were adjusted with Benjamini-Hochberg correction and significance was determined as FDR < 0.01.

STARR enrichment by TE class and lineage was calculated by grouping the overlaps by class or lineage of the TE that the peak summit overlapped and calculating the enrichment against the randomly shuffled peak summits as described above. TE lineage of origin data was obtained from ref. [65].

### ATAC-seq, ChIP-seq, TT-seq/GRO-seq read alignment to MER11B/LTR12C consensus sequences

For GP5d, ATAC-seq, TFAP2A ChIP-seq and TT-seq data were first aligned to the reference genome. Unique reads mapped to 92 MER11B elements that overlapped with STARR-seq peaks were extracted and mapped to the MER11B consensus sequence to obtain compiled reads.

For HepG2, ATAC-seq, NFYA ChIP-seq and GRO-seq data were first aligned to the reference genome. Unique reads mapped to 597 LTR12C elements that overlapped with HepG2 STARR-seq peaks were extracted and mapped to LTR12C consensus sequence to obtain compiled reads. Bwtools v.1.0 (bwtools aggregate) was used to plot compiled signal for ATAC-seq, ChIP-seq and TT-seq/GRO-seq at LTR consensus sequences[92].

### NucleoATAC analysis

NucleoATAC v.0.3.4 was used to predict the nucleosome positions by using ATAC-seq data[31]. Nucleosome occupancy signal around GP5d STARR-seq peaks was obtained by using NucleoATAC run function. Smoothed and normalized cross-correlation NucleoATAC signal was used to plot the nucleosome occupancy around the GP5d STARR-seq peaks.

### DepMap TFAP2A dependency analysis

DepMap CRISPRi screening data for TFAP2A was downloaded for colorectal and liver cell lines. CRISPR Chronos score (TFAP2A gene effect) and TFAP2A expression (Log2 (TPM+1)) for colorectal and liver cells was plotted by using GraphPad Prism 9[52].

### Motif analyses

All motif enrichments for each TE subfamily or STARR cluster were analyzed with AME from the MEME suite v. 5.0.2 with shuffled sequences as the background (ame --control --shuffle--)[93]. JASPAR 2022 CORE non-redundant vertebrate motif annotations were used as the motif file input. Motif clustering data was downloaded from (https://resources.altius.org/~jvierstra/projects/motif-clustering-v2.0beta/)[26]. Output from AME was read into R and the E-values were –log10-transformed. Each motif hit from AME analysis was assigned to a corresponding cluster. For each cluster, minimum E-value from all individual motif hits was selected, the columns were scaled with the R scale function (center = F) and plotted.

### Cluster analysis

In GP5d, RPKM normalized bigwig files for STARR-seq, ATAC-seq, ChIP-seq for H3K4me1, H3K4me3, H3K9me3, H3K27ac, H3K27me3, H3K36me3, p53 and 5-fluorouracil treated p53 and unnormalized CpG methylation bigwig from NaNoMe-seq were used to create a matrix file with deepTools v.3.5.0[75] with a region of 3kb in both directions around STARR-seq peak summits as the center reference point. (computeMatrix reference-point --referencePoint center – missingDataAsZero -a 3000 -b 3000). The resulting deepTools matrix was loaded into R and columns were centered with the R scale function. R kmeans function was then used to cluster the matrix with centers from two to nine and options nstart = 50, iter.max = 15. The resulting output was analyzed by elbow plotting and five clusters were determined to be the optimal number of clusters.

### Activity-by-contact model analysis

Candidate regions for the ABC model[38] were created using GP5d ATAC-seq data (see ATAC-seq analysis section for processing details) with the makeCandidateRegions.py with the ENCODE blacklist as excluded regions, 250bp peak extension and 150000 selected peaks (---regions_blocklist ENCFF356LFX.bed, --peakExtendFromSummit 250, --nStrongestPeaks 150000). Enhancer activity was quantified using the run.neighborhoods.py script with ATAC-seq and H3K27ac ChIP-seq (see ChIP-seq analysis section for processing details) and the expression table of TPM counts from Salmon analysis (see RNA sequencing section) and GENCODE v36 gene annotations as inputs.

Contacts were predicted with the predict.py script with the output from the previous steps with threshold set as .02, hic-resolution set at 5000, powerlaw scaling and GP5d H3K27ac Hi-ChIP data set as the contact mapping input (see Hi-ChIP analysis section for processing details) (--scale_hic_using_powerlaw, --threshold .02, --hic_resolution 5000).

### Statistical analysis and plots

All statistical and downstream analyses were performed in R v.4.1.2[94] Profile plots were created from the bigwig files with the R package soGGi v.1.24.1 using the regionPlot function with the option normalize = T[95]. The signal was smoothed with rollmean from Zoo package v.1.8-10[96]. Genomic annotation for STARR-seq peaks was performed with ChIPseeker[97]. All plotting was performed in ggplot2 v.3.3.6 from the Tidyverse suite v.1.3.1[98] except for the motif enrichments that were plotted with ComplexHeatmap v.2.8.0 [99] and enrichment heatmaps that were plotted with EnrichedHeatmap v.1.22.0[100], with code adapted from[45]. Illustrations were created with BioRender.com. Heatmaps in figure 3 and 4 were plotted by using deepTools[75].

## Data availability

Data generated in this study has been deposited into GEO under accession GSE221053. The publicly available data was accessed as follows:

GP5d and HepG2 STARR-seq data from GEO accession GSE180158. For GP5d, genomic and p53-null STARR-seq data from GSM5454433 and GSM5454435, and for HepG2 genomic STARR replicates 1 and 2 from GSM5454437 and GSM5454438, respectively.

GP5d ChIP-seq data for H3K27ac (GSM5454417), H3K9me3 (GSM5454420), H3K27me3 (GSM5454428), 5-FU treated mIgG (GSM5454414), untreated p53 (GSM5454412) and 5-FU treated p53 (GSM5454413) were acquired from the same study. GP5d H3K4me1 (GSM1240814) was obtained with the GEO accession GSE51234.

HepG2 ChIP-seq data for H3K27ac, H3K27me3, H3K36me3, H3K4me1, H3K4me3, H3K9me3, p53, NFYA and ZBTB33 replicate 1 and 2 from ENCODE (https://www.encodeproject.org/) with fastq file accessions ENCFF000BFD, ENCFF001FLQ, ENCFF001FMA, ENCFF000BEX, ENCFF901NZE, ENCFF000BFK, ENCFF257UIJ,ENCFF081VHA, ENCFF204TCE and ENCFF745CTV respectively. Replicate 1 for HepG2 ATAC-seq was downloaded with accessions ENCFF664UPL and ENCFF289UIB for read file 1 and 2, respectively.

A gene annotation GTF file was acquired from Gencode Release 36 for the reference chromosomes (https://ftp.ebi.ac.uk/pub/databases/gencode/Gencode_human/release_36/gencode.v36.annotation.gtf.gz).

A repeatMasker.txt (2021-09-03) file was downloaded from the UCSC table browser (https://genome.ucsc.edu/cgi-bin/hgTables). MER11B and LTR12C consensus sequences were acquired from RepBase (https://www.girinst.org/repbase/update/index.html).

GRCh38 chromosome sizes file (2020-03-13) file was downloaded from UCSC (https://hgdownload-test.gi.ucsc.edu/goldenPath/hg38/bigZips/latest/). Unified GRCh38 blacklist BED file (ENCFF356LFX, release 2020-05-05) was downloaded from ENCODE project (https://www.encodeproject.org/).

Transcription factor motifs were acquired from JASPAR 2022 CORE non-redundant vertebrate annotations (https://jaspar.genereg.net/download/data/2022/CORE). The position weight matrices in MEME format were used for downstream motif analyses. Motif clustering data was downloaded from (https://resources.altius.org/~jvierstra/projects/motif-clustering-v2.0beta/).

## Code availability

Custom code will be made available upon request.

## Acknowledgements

We thank HiLIFE research infrastructures including the Biomedicum Functional Genomics Unit (FuGU) and the FIMM NGS Genomics laboratory at the University of Helsinki. We thank the Center for Scientific Computing (CSC), Finland, for the computational infrastructure, Professor Lauri Aaltonen’s laboratory facilities for genomics work. We would also like to thank Justyna Kolakowska and Inga-Lill Åberg for their technical help with nanopore sequencing. BS was supported by the Academy of Finland (274555, 317807), Finnish Cancer Foundation, Sigrid Jusélius Foundation, and Jane and Aatos Erkko Foundation.

## Author Contributions

BS conceived and supervised the study and performed STARR-seq experiment with methylated library and TFAP2A ChIP-seq. KK performed STARR-seq and genomics data analysis together with DP who performed the ChIP-seq, TT-seq and GRO-seq analysis and epigenetic inhibition experiments. JX performed the NaNOMe-seq and Hi-ChIP and LF performed the ATAC-seq. KP helped with initial processing of NaNOME-seq data with the support from LA. BS, KK and DP wrote the manuscript with contributions from all authors.

